# A multi-host mechanistic model of African swine fever emergence and control in Romania

**DOI:** 10.1101/2024.01.09.574784

**Authors:** Brandon H. Hayes, Timothée Vergne, Nicolas Rose, Cristian Mortasivu, Mathieu Andraud

**Author notes:** Brandon H. Hayes **Email:**. **Author Contributions:** B.H.H., T.V, N.R., and M.A. designed research; B.H.H. performed research; B.H.H., T.V., N.R., and M.A. analyzed data; B.H.H. wrote the paper and T.V, N.R., C.M., and M.A. reviewed the manuscript. **Competing Interest Statement:** The authors have no competing interests to disclose.

## Abstract

African swine fever (ASF) has devastating effects on swine production, farmer livelihood, animal welfare, and biodiversity. Extremely difficult to control, epidemic management is further complicated when spillover between domestic pig and wild boar populations is suspected. To quantify ASF viral transmission between domestic pigs and wild boar, a spatially-explicit stochastic mechanistic model was constructed using village centroids to represent aggregated backyard pig herds and a hexagonal raster of forest coverage to represent wild boar abundance. The model was parameterized to the initial six months of the ongoing Romanian epidemic through approximate Bayesian computation. It was estimated that a median of 69.4% (interquartile range: 53.0–80.0%) of domestic pig herd cases came from other infected domestic pig herds while 20.4% (11.2–33.8%) originated from infected wild boar sources, and 8.4% (4.7–14.2%) stemmed from external sources not explicitly represented. Also, 31.9% of infected wild boar habitat (16.7–56.2%) originated from domestic pig herds and 68.1% (43.8–83.3%) came from neighboring infected wild boar populations. Furthermore, it was found that habitats with a forest coverage greater than 15% were 2.6 times more infectious and 5.3 times more susceptible than other habitats. All alternative control scenarios, including culling domestic pig herds upon local domestic pig or wild boar case detection, improved epidemic outcomes, with the greatest decrease in final epidemic size being observed from the reactive culling of entire villages following case detection. These results can be used to further inform policy recommendations in ASF-epidemic regions.

**Significance Statement:** The current African swine fever (ASF) pandemic is devastating to affected nations, and quantifying transmission parameters is critical to informing control strategies. Disease spillover between wild and domestic hosts further complicates control efforts, yet the influence of spillover events on epidemic propagation remains unknown. Using the context of Romania—one of the European nations with the most severe epidemic and where spillover transmission is strongly suspected—we show that targeting spillover mechanisms is critical for achieving holistic disease control, and then demonstrate the impact of alternative control scenarios had they been enacted. These results can inform control strategy policy decisions in the many nations at-risk for or actively experiencing ASF epidemics.

## Introduction

African swine fever (ASF), one of the highest consequence diseases of domestic pigs, continues to pose a grave threat to the global swine industry (1). Virus introduction and transmission pathways vary between affected areas, with some nations experiencing cases predominantly or exclusively among wild boar (e.g. the Baltic countries, the Republic of Korea), and others seeing cases mostly among domestic pigs (e.g. Romania, Ukraine, China, and much of southeast Asia) with likely spillover to-and-from wild boar (2–7). In many countries, spillover is highly suspected but the current surveillance bias is likely to prevent accurate evaluation of this transmission route (6). With no available treatment and an effective vaccine still out of reach, interrupting disease transmission is reliant on control strategies targeting these pathways (8, 9). Understanding the dynamics unique to each epidemic provides the best chance of achieving epidemic control (10).

Mechanistic models—which allow the quantification of transmission parameters, the prediction of epidemic trajectories, and the evaluation of the effectiveness of control strategies—are a proven way of providing quantitative information to guide potential control measures (11–14). However, current ASF models at the interface of domestic pigs and wild boar are rare and are not parameterized on real outbreak data, despite these events being suspected to play an important role in the propagation of some ASF epidemics (15). Understanding the transmission dynamics at this interface is necessary to design tailored control measures, adapted to the specificities of domestic pig rearing in regards of societal and economical aspects.

Since 2018, Romania has been facing an ASF epidemic of unprecedented scale, affecting both wild boar and domestic pigs (3). Genotyping has confirmed that the circulating genotype II African swine fever virus (ASFV) strain is identical to isolates both from EU member states (Lithuania and Poland) and Caucasus nations (16). The ubiquity of backyard pig farming in villages—with the majority of families having one or more pigs kept in low-biosecurity backyard holdings—provides an environment highly amenable to transmission between wild and domestic hosts (17). Understanding the dynamics of the initial conditions of the current sanitary situation could prove beneficial in improving control paradigms, especially for nations that have yet to experience an ASF incursion.

Using a multi-host spatiotemporal model of ASF transmission calibrated to the observed dynamics of ASF emergence in Romania, we aimed at estimating the relative contribution of domestic pigs and wild boar to overall epidemic propagation, quantifying parameters needed to explain the observed epidemic trajectory, and evaluating potential outcomes from alternative control strategies.

## Results

### Model fitting and inference

Frequency-dependent herd-to-herd transmission, density-dependent cell-to-herd transmission, second-order cell-to-cell transmission, and considering only forested cells as more suitable wild boar habitat (i.e. not also including neighbors of forested cells as more suitable wild boar habitat) were associated with the lowest MSE (Fig. S1, supplementary materials). Through observing trends in model fit, some formulations are seen to be systematically more supported by the data than others (e.g. all models that used density-dependent distance kernels or frequency-dependent step functions performed better than those that used a density-dependent step function).

The best fitting model’s prediction intervals satisfactorily captured the weekly incidence for each host (Fig. 1). When faceted spatially by six counties, the spatial distribution of the epidemic is further replicated in the majority of areas. The incidence of wild boar cases was captured in the weekly 95% prediction intervals for all counties, and only among domestic pig units in two counties—Braila and Tulcea—was a spike in weekly incidence in the observed data not replicated (Fig. S2, supplementary materials).

**Fig. 1.**
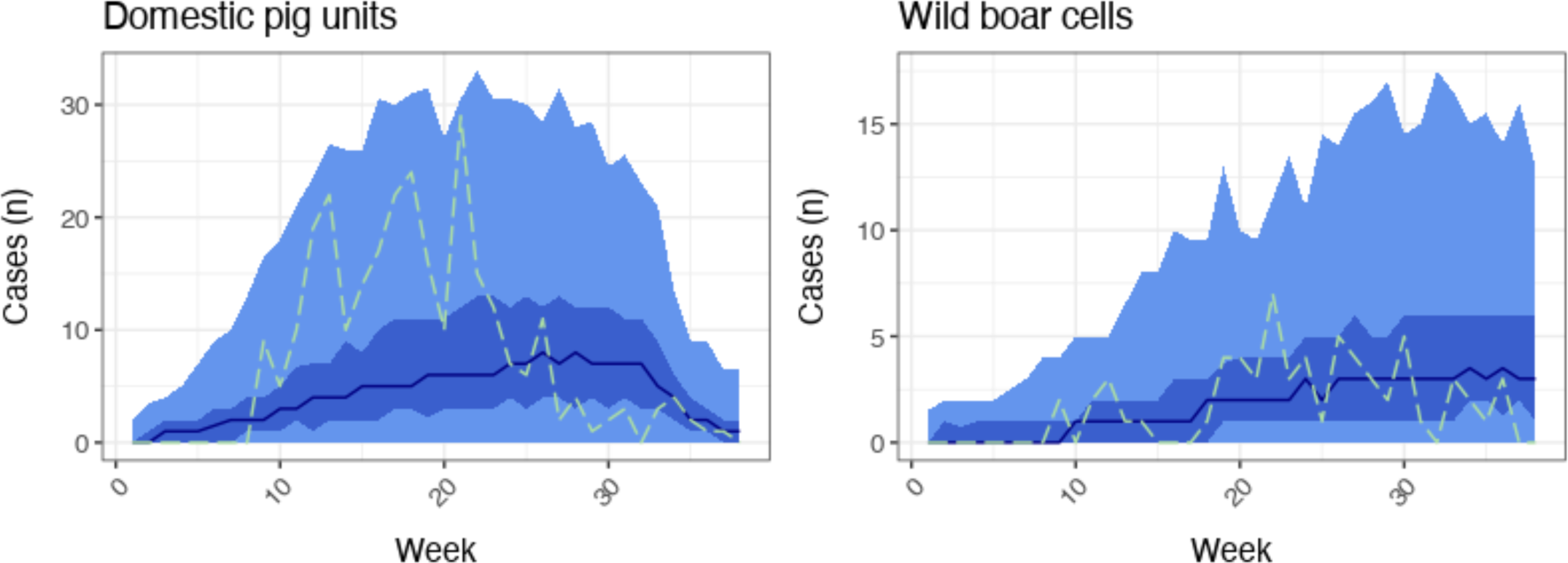
Epidemiologic trajectory of the best-fit model by epidemiologic unit showing captured dynamics. The light blue ribbon indicates the 99% credible interval—that is, a 99% probability the true epidemic estimate lies within this region—while the darker blue ribbon indicates the 50% credible interval, the darkest blue line the median of 1000 simulations, and the dashed green line the trajectory of the observed epidemic.

Nine parameters were estimated in the best-fitting model: five transmission rates between unit types (including transmission from external sources of infection to domestic pig units), the relative infectivity and susceptibility of wild boar cells based on habitat quality, and the relative infectivity and susceptibility of villages during the winter holiday slaughter period (Table 1, main text, Fig. S3, supplementary materials). Though the distance kernel (δ) for transmission between domestic pig units was estimated as well, the best fitting model utilized a step function (with the force of infection constant at distances shorter than 20 km and null otherwise) to scale the force of infection between domestic units with increasing distance. We estimate that a susceptible village was infected by an infectious village a median of every three weeks, while a susceptible wild boar habitat cell was infected by a neighbouring infectious cell every four weeks. Τhe transmission rate from an infected cell to village herds was estimated at approximately two cases every three weeks, and the transmission rate from village herds to the local wild boar environment was estimated at approximately one case per week.

**Table 1.** Best-fitting model parameter values (with credible intervals for estimated parameters)

| Parameter | Description | Value (95% CrI)* | Source value | Source |
| --- | --- | --- | --- | --- |
| $\beta_{i,j}$ | Transmission rate between herds (week<sup>-1</sup>) | 0.307<br>(0.0572–0.578) | NA | Estimated |
| $\beta_{i,q}$ | Transmission rate from cells to herds (week<sup>-1</sup>) | 0.670<br>(0.0177–1.86) | NA | Estimated |
| $\beta_{p,j}$ | Transmission rate from herds to cells (week<sup>-1</sup>) | 1.02<br>(0.131–1.92) | NA | Estimated |
| $\beta_{p,q}$ | Transmission rate between cells (week<sup>-1</sup>) | 0.281<br>(0.0230–1.47) | NA | Estimated |
| $\beta_{ext}$ | Transmission rate from external sources to herds (week<sup>-1</sup>) | 5.15E-4<br>(3.29E-5–9.69E-4) | NA | Estimated |
| $\psi_p$ | Relative infectivity of cells with inadequate forest coverage | 0.391<br>(0.0190–0.939) | NA | Estimated |
| $\phi_q$ | Relative susceptibility of cells with inadequate forest coverage | 0.187<br>(0.00514–0.818) | NA | Estimated |
| $\psi_{vil}$ | Relative infectivity of villages during holiday slaughter period | 0.230<br>(0.0162–0.470) | NA | Estimated |
| $\phi_{vil}$ | Relative susceptibility of villages during holiday slaughter period | 0.213<br>(0.0180–0.482) | NA | Estimated |
| $\sigma_i$ | Detection rate for infected herds (week<sup>-1</sup>) | 0.33 | 0.25–0.5 | (81) |
| $\sigma_p$ | Detection rate for infected cells (week<sup>-1</sup>) | 0.125 | 0.125–0.225 | (82) |
| $\gamma_{i.vil}$ | Recovery rate for infected villages (week<sup>-1</sup>) | Variable by county and time period** | NA | Observed |
| $\gamma_{i.ind}$ | Recovery rate for infected industrial sites (week<sup>-1</sup>) | 1 | NA | Observed |

\*95% credible interval
\*\*See table S1 for spatiotemporally-explicit village recovery rates

Using the best-fitting model that captured the observed epidemic trajectory, the contributions of different epidemiological unit classes were estimated (Fig. 2). Among domestic pig herds, a median of 68.7% (IQR 52.4–80.6%) of infections came from other pig herds while 21.3% (11.8–32.3%) came from neighboring infected wild boar populations. Conversely, among wild boar cells, 67.7% (45.4–83.7%) of infections came from other wild boar cells while 32.3% (16.3–54.6%) came from domestic pig units. The external force of infection, representing the mean of all additional forces of infection not otherwise captured, accounted for very few infections to domestic pig units (8.69%, 4.62–15.3%). Furthermore, it was found that habitats with a forest coverage greater than 15% were 2.6 times more infectious and 5.3 times more susceptible than habitats with insufficient forest coverage.

**Fig. 2.**
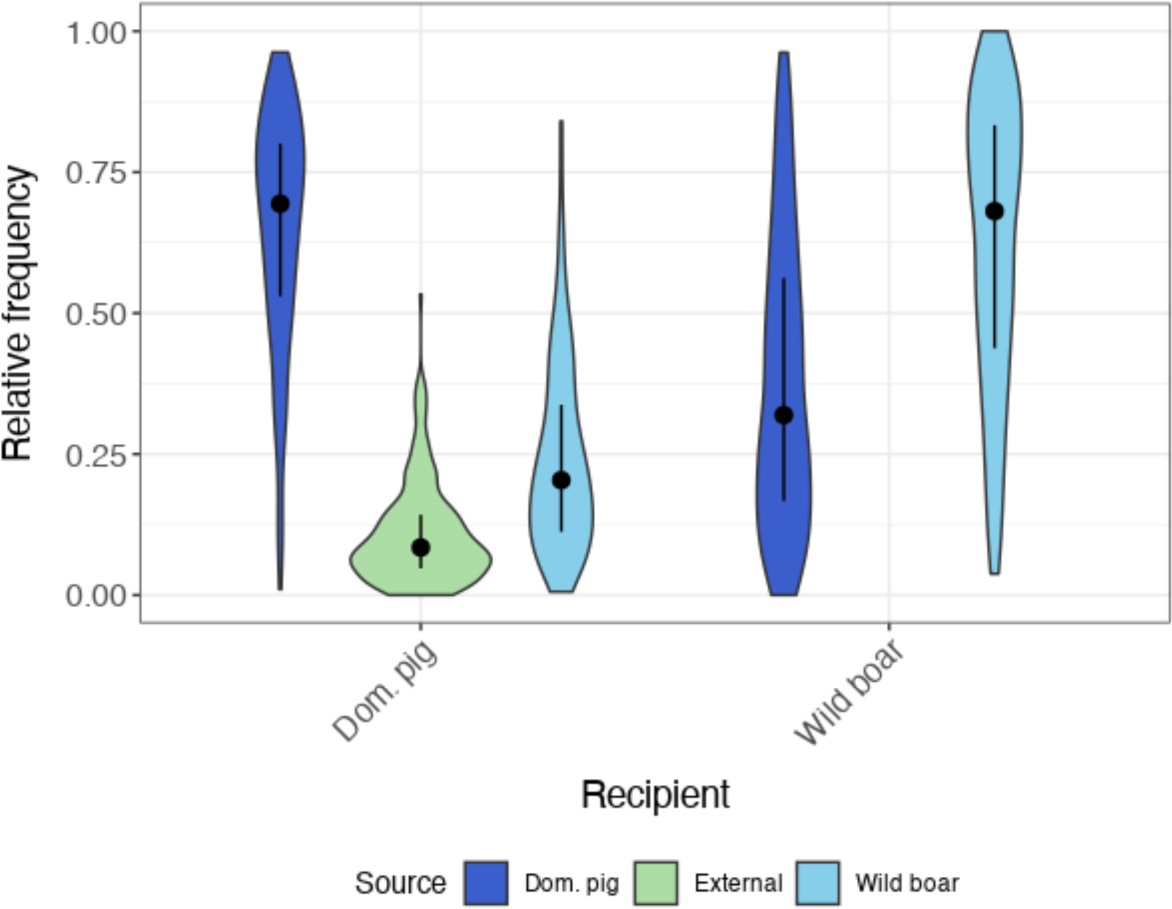
Relative contribution of domestic pig units and wild boar cells to epidemic propagation, as estimated in the best-fit model from 1000 simulations. For each recipient (as indicated on the x-axis), the y-axis indicates the relative frequency of infection for the color-coded given source of infection, with the width of the violin plot demonstrating the frequency of the value. The black dot inside the violin plots indicates median value with the bar indicating the interquartile range (IQR). For domestic pigs, a median of 69.4% (IQR 53.0–80.0%) of infections can from other domestic pig units, 20.4% (11.2–33.8%) came from wild boar cells, and 8.43% (4.73–14.2%) came from other sources not explicitly represented (via the external force of infection). Among wild boar cells, 31.9% IQR 16.7–56.2%) of infections came from domestic pig units, while 68.1% (43.8–83.3%) of infections were from other cells.

### Alternative management scenarios

Alternative control strategies were chosen to consider improvements in either the timeliness of case detection in domestic pigs or outbreak management in both domestic pigs and wild boar. They were evaluated through the end effects on host-specific final epidemic size and relative host contribution. Improvements in passive surveillance among domestic pig units, alternative culling strategies including both preventive and reactive culling of domestic pig units, and environmental sanitation through wild boar carcass removal were all examined. Different control strategies resulted in varying reductions in epidemic size (Fig. 3) and relative host contributions (Fig. 4).

**Fig. 3.**
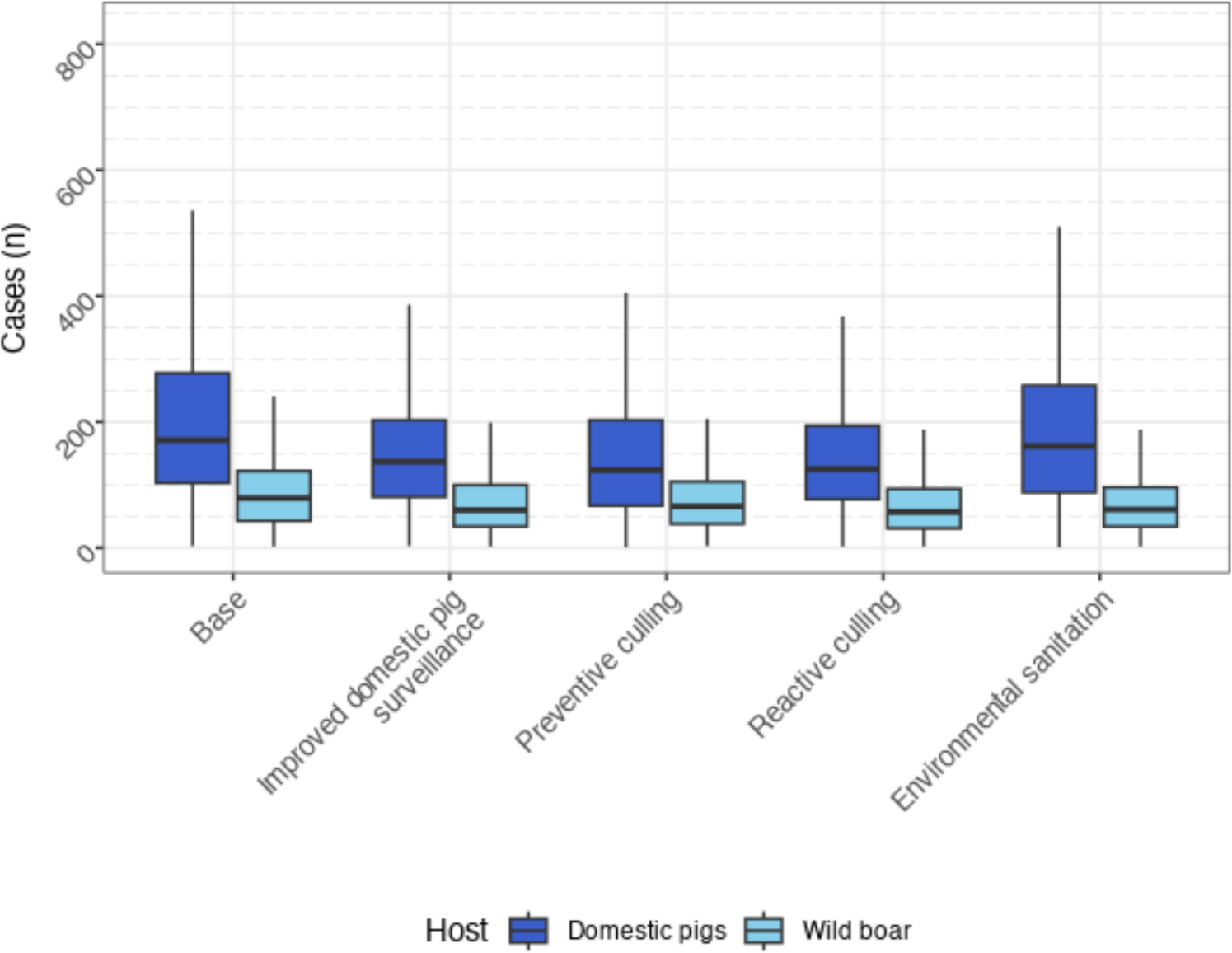
Effects of alternative control strategies on host-specific final epidemic size. Boxplots represent median and interquartile range (IQR), whiskers extend to 1.5 times the IQR, and dots represent outlier data points across 1000 simulations per scenario.

**Fig. 4.**
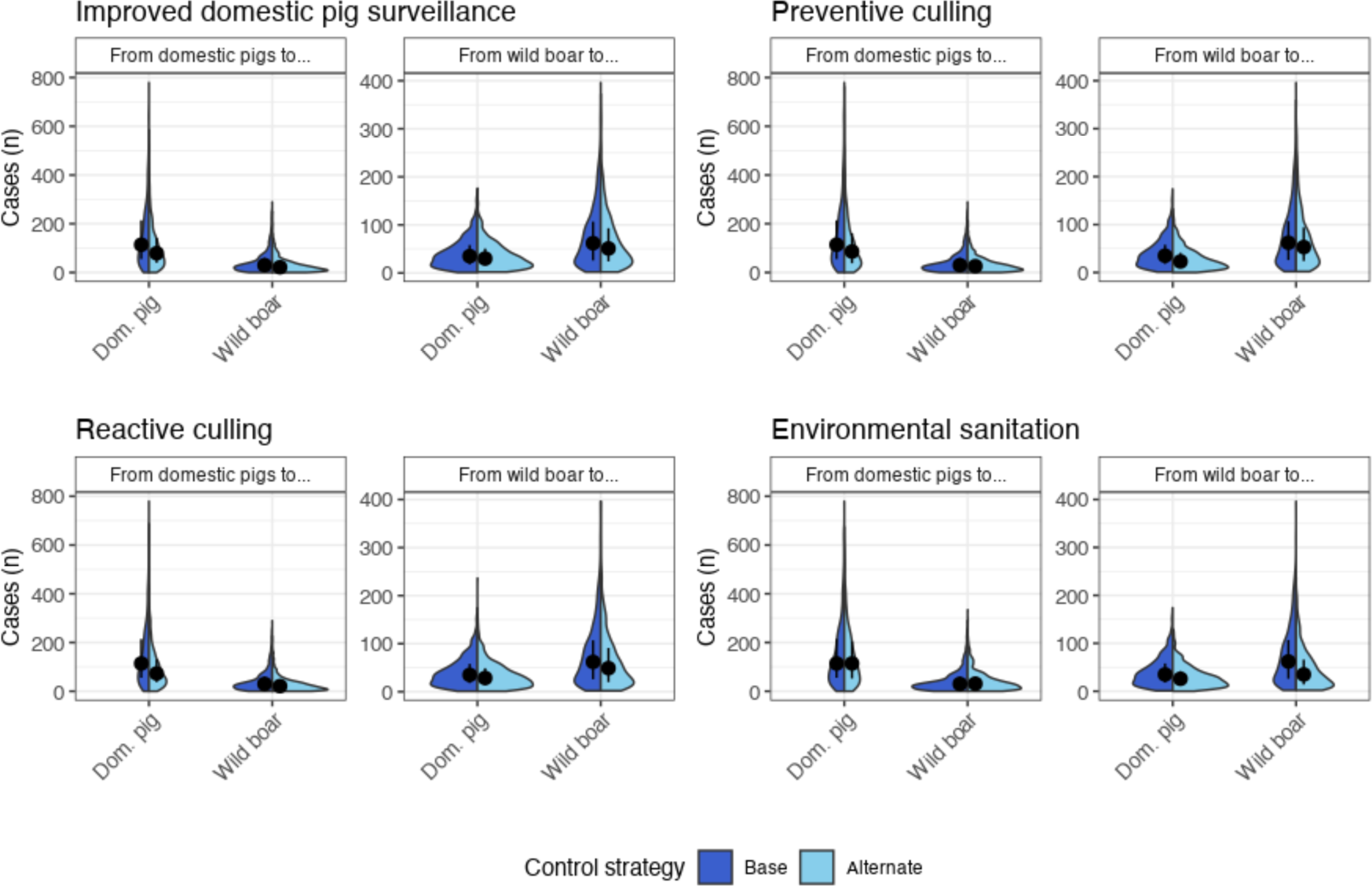
Effects of alternative control strategies on transmission frequency between hosts. Each pair of plots compares the absolute number of infections from each source to each recipient for the baseline scenario (dark blue) and the alternative control scenario (light blue) across 1000 simulations for each scenario. The black dot inside each split violin plot indicates median value with the bar indicating the interquartile range. Improving passive surveillance among domestic pig units was simulated through reducing the average duration of the undetected-infectious period from three to two weeks. Reactive culling of an entire village following case detection in that village was simulated through forcing an average infectious period duration of one week across all villages. Preventive culling of domestic pig units following detection of a nearby wild boar case was simulated by transiting domestic pig units in an infectious-detected cell to the R state upon cell case detection. Lastly, environmental sanitation through wild boar carcass clearance was simulated by having cells progress to the recovered state following a four-week infectious-detected period instead of remaining in the infectious state.

Improving passive surveillance among domestic pig units through reducing the average duration of the undetected-infectious period from three to two weeks resulted in a median of 23.3% (95% highest density interval (HDI): 18.8–27.4%) fewer domestic pig unit infections and 18.9% (14.2–23.5%) fewer wild boar cell infections, which translates to a disease-free area of almost 400 km^2^. These reductions in transmission frequency were seen primarily from between domestic pig units, though cross-host reductions were also present (Table 2).

**Table 2.**
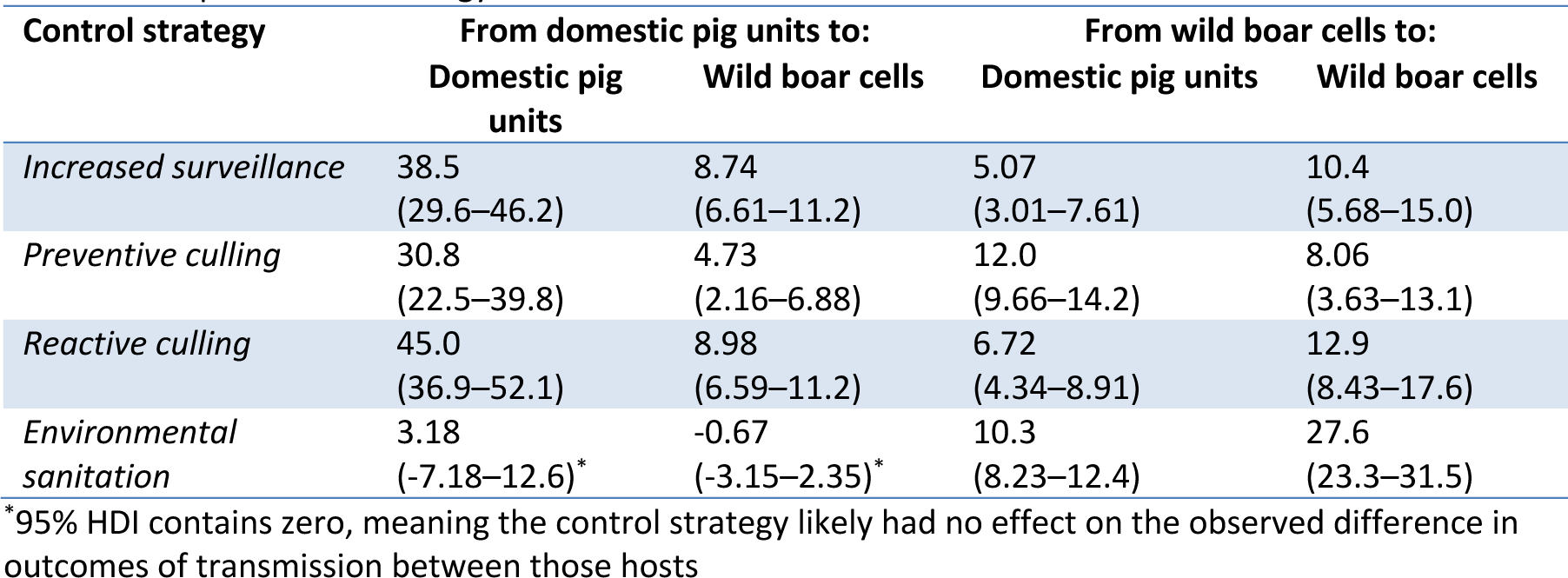
Median reductions in case numbers (with 95% highest density interval (HDI)) by route of transmission per control strategy.

Preventive culling of domestic pig units located within infected cells upon the detection of the first wild boar case resulted in a median of 28.6% (24.1–32.8%) fewer infections among domestic pig units and 13.9% (8.8—19.1%) fewer infections among wild boar cells, with most reductions in overall epidemic size resulting from decreases in transmission between domestic pig units. Significant decreases in transmission from wild boar cells to domestic pig units were also evident.

Reducing the duration of the village infectious period from the value observed in the epidemic (range: 1.3– 2.9 weeks) to 1 week to mimic reactive culling of all pigs from an entire village upon case detection in that village resulted in a median of 28.5% (24.5–32.0%) fewer infections among domestic pig units, and 24.5% (20.0–28.8%) fewer infections among wild boar cells. In addition to the large reduction in infections originating from domestic pig units, an almost 30% reduction in the number of infections between wild boar cells was also observed.

Environmental sanitation through wild boar carcass clearance was simulated by reducing the duration of the infectious period of wild boar cells to only for four weeks. This strategy resulted in a median of 7.4% (2.1–12.2%) fewer infections among domestic pig units and 20.8% (15.9–25.2%) fewer infections among wild boar cells. The largest relative decrease in transmission from this strategy was among infections from wild boar cells to other wild boar cells. Benefits of this strategy on decreasing the number of infections originating from domestic pig sources was not observed.

## Discussion

A mechanistic model of ASFV spread that simulates transmission both within and between domestic pig and wild boar host populations was constructed and parameterized using data from the on-going epidemic in Romania. Our analysis focused on the initial 30 week period of emergence, where the initialization and subsequent augmentation of control strategies led to non-uniform application of control measures both temporally over the study period and spatially between subnational divisions (counties) (18, 19). These differences were accounted for in the modeled transmission dynamics, and to do so our model included both spatial-dependence and seasonal-forcing of infectious period parameters in an attempt to reflect the varied and changing control pressures. The final result was a model where the overall trends observed in the epidemic emergence period were able to be replicated through stochastic simulation.

### Estimated parameters and replicated dynamics

Romanian ASF epidemic dynamics were captured by estimating transmission rates among and between villages and the environment, the latter of which was defined through estimated wild boar home ranges. Through disentangling transmission rates by the source and recipient hosts, and including unit subtypes with their own relative infectivity and susceptibility, the relative contribution of each epidemiological unit to epidemic propagation was able to be elucidated.

In the Russian Federation, the transmission rate of the current pandemic ASFV strain between pig farms was previously estimated at approximately 1–1.5 farms per week, whereas we estimate a median of one village infected by another village every 3 weeks (20). Though both transmission rates reflect local transmission patterns not related to industrial transport, our lower estimate is likely a reflection of differences in scale, with our model examining village herds en-masse as opposed to individual small-holder farms. Further estimates of the rate of transmission between domestic pig herds from other ASF epidemics would provide additional results for comparison.

Rates of wild boar spread of ASF have been estimated at 2.5 km/month in Poland (21), 2–5 km/month in Poland and the Baltic nations (2, 22), and 1.5 km/month in Poland (23). Given the 2.7 km apothem of our habitat hexagon, our estimated median rate of one 25 km^2^ patch every 4 weeks is in concurrence with the high-end of the median estimated velocities. Further, differences in transmission rates likely reflect geographic differences between areas of estimation.

A transmission rate from wild boar to domestic pigs has been previously estimated for the ASF situation in Poland (21), but it was derived from estimated contact rates between wild boar and domestic pig on free-roaming pig farms in savannah-like habitat in Spain (24). Conversely, contact patterns in England between wild boar and domestic pig farms were estimated at upwards of 7–14 contacts per month (25). Assuming a transmission probability from wild boar to domestic pigs of 0.167 (26), this equates to an expected transmission rate of 1 case every 1.7 to 3.4 weeks, in-line with our own estimates of two villages infected every three weeks by infectious wild boar cells.

Environment with sufficient wild boar habitat was estimated to be 2.6 times more infectious and 5.3 times more susceptible to ASF contamination than insufficiently-forested areas. Wild boar habitat preference for forested areas is well established, as is their non-preference for non-forested areas (27). Further, wild boar foraging behavior is understood to not be randomly distributed throughout forested areas, but is informed by resource proximity (28). Similarly, in our model, within forested habitat cells, additional factors like village edge size and proximity to nearest forest edges would likely inform the infectiousness and susceptibility of a forested area towards a village. Future studies at a finer resolution, incorporating two-dimensional representations of villages, could further elucidate the explicit factors behind such a pronounced increase in the infectiousness and susceptibility of forested areas.

In order to replicate the observed epidemic dynamics among villages, parameters to account for decreases in the overall domestic pig population from the mass nationwide culls during the holiday slaughter season had to be included in the model. Local customs play heavily into ASF transmission, with pig rearing commencing in the summer months in preparation for the December festivals, and the winter holiday slaughter acting as a strong external influence on ASF infection and propagation between villages (19). Our modeling outputs suggested that the infectivity and susceptibility of villages after the start of the holiday slaughter season underwent a median reduction in infectiousness and susceptibility of 77% and 79%, respectively, as compared to before the holiday season. This large decrease, which was necessary to allow epidemic extinction in the simulations as had occurred in the observed scenario, is most likely the consequence of decreases in the overall domestic pig population from the mass culls due to traditional festivities. Indeed, the large production of pigs from July to December substantially increases the susceptible swine population over that period, and half of the total annual slaughter of pigs in the country occurs in the month of December (29). Research into the social dynamics around pig farming during this time period would help to further clarify the accuracy of this fitted value. As the subsequent epidemic wave in 2019 was initiated at a similar period, it is possible that this temporal consistency is associated with this annual social dynamic, though further investigation is required to confirm this hypothesis.

Approximately 21% of domestic pig infections were estimated to come from wild boar sources, and roughly 32% of wild boar infections were estimated to originate from domestic pig sources. This transmission estimate from wild boar to domestic pigs is in-line with estimates from South Korea, where it was shown that six out of 14 farms had approximately a 60% risk of the main source of infection being wild boar, equating to a frequency of 26% (30). Such a degree of spillover suggests that measures acting on one population could destabilise the whole system. This dynamic could be of practical importance especially in the case of vaccination. Deciding to whom a vaccine should be delivered, either targeting domestic pigs or considering an oral vaccine to be introduced to wild boar, should be further explored to improve control efforts in both hosts (31).

Without transmission from domestic pigs to wild boar, the spatial dynamics of the observed wild boar epidemic were unable to be replicated. In those simulations, wild boar cases occurred solely in Tulcea county (the county of initial spread) over the examined period, suggesting the essential role that between-host transmission played in the observed epidemic (Fig. S4, supplementary materials). Further confirmation of the role of interhost transmission in epidemic propagation can be achieved through phylodynamic approaches, where phylogenetics is coupled with epidemiological data. Such an approach demonstrates particular promise at elucidating transmission parameters and quantifying spillover events (32). However, the choice of use of such methods has dependence on viral genetic characteristics, and ASFV, being a large double-stranded DNA virus—therefore subject to very slow mutation rates—may currently limit successes of this approach. Deciding upon whole genome or single-gene approaches, along with identifying if genetic variability can be captured with just a small number of sequences among outbreaks, are current hurdles with such a method (33, 34).

### Alternative management scenarios

Following determination of the relative host contributions to the overall epidemic, alternate management scenarios were explored. All examined alternative control scenarios resulted in reductions in final epidemic size, though not through equal reductions among the different transmission routes. Strategies that employed village-wide culling—either preventively upon nearby wild boar case detection or reactively following case detection in that village—resulted in the greatest decreases in epidemic size among domestic pig herds. Unfortunately, reactive culling of herds neighboring infected herds within a village was reported to be difficult to justify to small-scale Romanian farmers, and possibly led to illegal movement activities generating counter-productive effects (18). In contrast, during the ASF outbreak in the Republic of Korea in September 2019, reactive culling of all swine herds within a 3 km radius of an infected herd was successfully instituted, resulting in complete outbreak control among domestic pigs in under four weeks (35). There are likely multiple cultural, social, and political factors behind these differences in the acceptance of such a strategy, and decision makers would benefit from further social science research into them.

In addition to reactive culling of pig herds, preventive culling following nearby wild boar case detection was similarly effective at decreasing final epidemic size. Field research has shown wildlife contact to be an important contributor to cases among domestic pigs. Phylogenetic results in Poland revealed that while separate genetically-distinct ASFV incursions were responsible for clusters of outbreaks, within those clusters wild boar were suspected to be responsible for transmission to domestic pig herds (36). In Estonia, direct contact with wild boar was suspected to be the inciting cause of outbreaks on two out of 28 investigated farms (37). Beyond only direct transmission, a scoping review of articles on wild-domestic ASFV transmission concluded that when such between-host transmission occurs, indirect routes may in fact be more common (38). Our results are in accordance with one of the models out of the ASF Modelling Challenge, where preventive culling of domestic pig herds nearby to detected wild boar cases resulted in a decrease of almost 20% in the final number of infected pig herds (39). Acceptance of such a strategy by farmers could be difficult without targeted educational programs. However, as highlighted in (39), the cost-benefit ratio of such a strategy would have to be established given the presumed substantial costs.

Reducing the undetected-infectious period of villages from three to two weeks—as can occur through improved case reporting—reduced final epidemic size for both the domestic and wild compartments and resulted in fewer transmission events in all directions. Through providing educational opportunities that lead to better awareness of symptoms and risks, pig holders could be more sensitive to the risk of transmission to others, and could react more rapidly. Indeed, the risk of transmission due to individual farmer behavior during emergency sales has been shown to be high (40). In the observed data, a progressive decrease in the estimated quarterly infectious periods is observed, which could be attributable to increasing awareness by pig holders over the course of the epidemic.

Environmental sanitation—as would occur through wild boar carcass clearance and simulated through limiting the infectious period of wild boar habitat cells—had the least effect on mitigating domestic pig outbreaks. Given the smaller proportion of infection events from contaminated environment to domestic pigs, this was expected. However, carcass removal has been shown to be a critical supportive intervention against ASF, and is true especially when instituted during epidemic onset; as seen in the successful elimination of ASF in both Belgium and the Czech Republic (41–43). Previous study indicated that at least 50% of carcasses must be removed for this strategy to have an effect, so it can be assumed that, in our model, sanitation of an infected cell would be defined as the capture of at least 50% of the carcasses in a 25 km^2^ area (44). While this could be perceived as an unrealistic task given that carcass removal efforts in Latvia were estimated to detect less than 0.5 carcasses / 100 km^2^, our model assumes a uniform landscape (outside of forest coverage) that is free from any within-cell fragmentation. Rivers, highways, commercial infrastructure and other geological features— which have all been shown to affect transmission dynamics— can all contribute to such landscape division thus limiting the overall area to search for carcasses (45).

Achieving control of an ASF outbreak is a complex task requiring the consideration of the effects of management decisions both within and between ecological compartments, and involving coordinated efforts between stakeholders from veterinary services, animal agriculture, conservation and hunting groups, law enforcement, and government, among others. In areas rich in backyard farming, as long as domestic pig herds continue to be a source of infection for wild boar and vice-versa, without the explicit targeting of achieving disease control in both compartments, updating epidemic control objectives to consider low levels of circulation in the wild compartment may be necessary.

### Model formulation

A recent review identified the milieu of approaches to estimating parameters and assessing control strategies of multi-host epidemics, though only five of the 56 examined approaches used a similar point-location-over-habitat-raster framework (46). Noteworthy, this approach was recently employed by a team in the ASF Modelling Challenge (47, 48). Alternative approaches, where wildlife is either explicitly individually simulated or—on the other end of the spectrum—represented through a single encompassing parameter such as an additional force of infection, can be either too computationally intensive or not granular enough for estimating the contribution of multiple routes of disease transmission to epidemic propagation in a multi-host environment.

The epidemic data that were used to calibrate the model supported frequency-dependent transmission between domestic pig herds, as occurs when contact rates between hosts scale independent of population density, bound by a step function at 20 km (10, 49). While other mechanistic ASF models used power-law transmission kernels that assumed maximum local transmission distances of 2 km (50–54), these models accounted for non-local transmission through network movements and in some cases were also tailored to industrialized populations (50–52). Without considering industrial networks, and to capture non-reported pig movements between villages which principally occur at a small scale, our maximum distance was increased accordingly. Transmission between villages was suspected to reflect local trade and social dynamics. While the fitted transmission formulation may be realistic regarding local spread between villages, this is less likely the case among industrial sites where transport distances of pigs can be many times greater. Additionally, infection transmission among the transit network is not being modelled, and the external force of infection exists to capture these long-distance jumps in transmission that—in the absence of movement records—are otherwise extremely difficult to model mechanistically.

In our wild boar model, forest coverage was used as a proxy variable for species abundance, selected for its availability and its established usage for representing wild boar (21, 47, 55–58). Habitat suitability models, where the suitability of a location is based on observed environmental relationships, are already extensively used in the field of ecology (59, 60). They are, however, not without a few key assumptions: that the species-environment relationship is already at equilibrium, and the data of the environmental variable being used is at an appropriate resolution to capture the species’ niche (59). Through including both forested and forest-adjacent cells as suitable habitat in our model, and using cells sized to previous estimates of wild boar group home ranges, we were able to account for the latter (27). Given the relatively short time span of our model, effects on population dynamics from natural variations (births and deaths), hunting, and disease-induced mortality were not considered, and the wild boar ecosystem was assumed to be at equilibrium over the period. Over longer periods, temporal changes in individual landscape variables, as is the case with seasonal land-use in agriculture or forestry exploitation on forest distributions, are important considerations in species representation. As was the case with ASF, recent research revealed that artificial landscape fragmentation was efficient at slowing the spread of ASF among wild boar (45). Future ASF models at the domestic-wildlife interface may want to consider combinations of habitat suitability variables, or use more advanced ecological models (e.g. friction models) for accurately representing wildlife hosts, especially across seasons and at larger time scales than our model. However, the additional costs in complexity and computing time would have to be considered prior to such a decision.

The dynamics seen with the current genotype II strain revealed the ability of the virus to persist environmentally without repeated re-introduction events (61). Both low-density endemicity and ASFV propagation have been observed and attributed to combinations of ASFV environmental stability, carcass-mediated transmission, and wild boar behavior, though viral shedding from persistently-infectious survivors has also been proposed (62). When ASF enters the local wild boar population, a large and rapid decrease in population (approximately 85% of all individuals) is first observed (63). Following this population crash, ASF can become endemic with a low prevalence—as currently is suspected in northern Europe—and continue to be transmitted through direct and carcass-based transmission (44). A similar population crash could be expected with the introduction of ASFV into the wild boar-dense regions of Romania, which would then require frequent transmission from domestic pig herds to result in the observed dissemination of cases in our area of study. Indeed, underreporting among wild boar cases is a known limitation of the surveillance efforts during the six-month period of our investigation. Further informed surveillance will allow a greater understanding of wild boar case distribution and spatial correlation.

As our model did not include internal infection dynamics within each village, many of the nuances that would contribute to reinfection were unlikely to be captured through a single village-wide force of infection parameter (64). Neighbors of ASF-infected farms slaughtering and storing their pigs prior to investigation to avoid culling of their herds (as reported by local investigators), as well as the continued practice of swill feeding, contribute to village reinfection (64). Conversely, the limited duration of the study period also is likely to have precluded the full capture of suspected reinfection events. Though the absence of a reinfection mechanism is not fully representative of the observed data, the overall significance of this assumption was considered low given the low percentage of villages that were affected more than once. In contrast, wild boar cells were assumed to remain perpetually infectious once infected; as mentioned previously environmental elimination is highly improbable without more than 50% of infectious carcasses being discovered (44). As wild boar surveillance was imperfect and variable during this initial epidemic period, it is unlikely that more than half of all infectious carcasses were discovered. This is commensurate with the results of initial selection of model infection states, where the best-fitting models were those that considered perpetual environmental contamination (see supplementary material: infection state selection).

### Model and data limitations

Formulating and fitting a model to an active epidemic presents unique challenges and limitations. In capturing outbreak trends, two observed spikes in incidence among domestic pig were not able to be replicated in our model, suggesting the influence of additional local dynamics not explicitly considered. Incorporating village-specific demographic factors or within-village dynamics could potentially improve the ability of the model to replicate these observations. Further, such internal dynamics would be important factors to consider prior to recommending enaction of the assessed high-consequence control strategies (i.e. large-scale culling). However, accounting for within-village transmission dynamics would require actual geolocation of domestic pig holdings—which is currently lacking—and obtaining such census data should be a consideration among Romanian stakeholders. An additional limitation in our model is the assumption that all infected cells were eventually detected. The true wild boar case count is likely even greater than that of domestic pigs, and the estimated transmission rates fitted to imperfect surveillance data may be biased. However, by fitting our model to infected environmental patches rather than individual wild boar cases, the suspected underreporting bias among wild boar was likely buffered. Examining additional time periods of the Romanian epidemic, following the initial phase in 2018 and when consistent surveillance efforts were established, could help to further refine estimated transmission rates.

### Policy impact and real-time support

Through using a static model of livestock and a simplified proxy for wildlife, a multi-host epidemiological model was able to be developed and employed on a (relatively) short time-scale: a pace sufficient to inform on-going outbreak scenarios. By substituting the current village locations with those of another region and regenerating the habitat raster for that area, the model can be rapidly adapted to additional outbreak locations or even predicted outbreak locations. Such flexibility, during times of regularly updated data and evolving control measures, could enable real-time policy support (47). This approach could prove beneficial for exploring control strategies of other high-consequence diseases at the livestock-wildlife interface in an emergency setting. Indeed, of the 53 livestock diseases in the United States of America that are reportable to WOAH, 42 have a wildlife component, with similar livestock-wildlife disease risk and susceptibility seen in Europe (65, 66). Extending the livestock component to also include commercial network trade would allow this type of model to be employed in areas with greater industrialized livestock populations.

Through the rapid deployment of a model of an ongoing outbreak, real-time policy support for animal health stakeholders can be a reality during the period of epidemic onset, a period whose dynamics are invariably different than those of other epidemic phases (67). The use of mathematical modeling to inform decision-making has become standard practice for public health, and with sufficient resource availability it can become a central tool in veterinary public health too (68).

## Materials and Methods

### Data

Four datasets were used in the construction and calibration of the model: outbreak records, village spatial data, industrial swine herd locations, and land cover to approximate wild boar distribution. Case records, which include the date of occurrence, coordinates, type of host (swine or wild boar) and associated farm type (backyard or industrial) were retrieved from the World Organization of Animal Health (WOAH) Animal Disease Information System database for the period from initial case detection (June 10, 2018) to the end of the first year (Dec. 31, 2018) (69). Cases were restricted to the southeastern region of Romania where the initial epidemic spread was witnessed—the counties of Braila, Calarasi, Constanta, Galati, Ialomita, and Tulcea. Village spatial data was retrieved from the Romanian National Agency for Mapping and Real Estate (ANCPI), providing a total of 986 villages in the six counties (70). Locations of industrial sites (n = 55) were retrieved from data available through the county-level directorates of the National Sanitary Veterinary and Food Safety Authority (71). Forest coverage was obtained from CORINE Land Cover satellite imagery (72).

### Model agents

A spatially-explicit model of the southeastern Romanian epidemiological system was constructed through layering an agent-based model of ASF transmission between domestic pig herds over a cellular automata model of ASF transmission in wild boar populations. The model explicitly allowed for disease transmission both within and between layers (Fig. S5). As individual backyard herds were not able to be explicitly located, and per conclusions from field investigations that recommended counting outbreaks in the form of infected and cleared villages, backyard herds were aggregated at the level of the village with each village acting as a single epidemiological unit (18). Industrial sites were also included in the domestic pig herd population, however being limited in number in our area of geographic interest (representing 55 out of 1041 herds) and being principally large integrative herds that likely play only a minor role in transmission to backyard herds, were not explicitly differentiated from villages. Village locations were represented by the centroid of each village. The wild boar population was represented through a rasterization of the southeast Romanian area of study of 25 km^2^ hexagons, sized to estimated wild boar home ranges (27). It is expected that wild boar abundance influences local disease transmission, so hexagons were characterized by their habitat quality which was defined through forest coverage (73). Using a threshold of 15% to define high and low forest coverage cells allowed for the assigning of different transmission parameter values (74, 75).

### Infection process

The infection state process for each model agent (domestic pig herd or wild boar cell) followed a modified SIR (susceptible (*S*), infectious (*I*), recovered (*R*)) framework, with the infectious state partitioned into infectious-undetected (*Iu*) and infectious-detected (*Id*) states. Latency was not considered as the model runs in discrete weekly timesteps and the mean incubation period for the ASFV genotype II strain of interest is frequently estimated at a week or less (76–78). Re-susceptibility was not included as only 13% of infected villages experienced a resurgence of ASF following outbreak clearance and augmentation of the domestic pig model through accounting for reinfection throughout the study area was not supported by the data (supplemental material: epidemic analysis). To be able to identify the contribution of the different transmission routes, five transmission rates (β), defined as the rates at which a susceptible agent acquires infection from an infectious agent per unit time, were considered: from infectious herd *i* to susceptible herd *j* (β_!,#_), from infectious cell *p* to a susceptible herd *j* (β_$,#_), from infectious herd *i* to susceptible cell *q* (β_!,%_), from infectious cell *p* to susceptible cell *q* (β_$,%_), and from external sources of infection not explicitly represented (including long-distance trade among the commercial network and reintroduction pressures from cross-border activity with neighboring infected countries) to susceptible herd *j* (β_&’(,#_). No external force of infection was included for wild boar cells as only local transmission was considered appropriate between wild boar.

At time *t*, the force of infection experienced by susceptible herd *j* was given as the sum of all forces of infection (λ) exerted by infected herds *i* and infected cell *p* that contains herd *j*, plus the external force of infection (Eq. 1). The force of infection experienced by susceptible cell *q* was given as the sum of all forces of infection exerted by infected herds *i* within susceptible cell *q,* and neighboring infected cells *p* (Eq. 2). The total force of infection on the susceptible unit (herd *j* or cell *q*) was further modified by its relative susceptibility (ϕ_*j*_ or ϕ_*q*_) determined by forest coverage for cells and temporally by season for domestic pig herds.

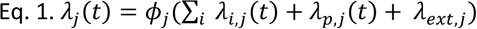

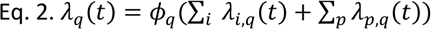

With λj being the total infectious pressure experienced by a susceptible herd and λq the total infectious pressure experienced by a susceptible cell.

Individual forces of infection (Eq. 3–7)—from either infected herd *i*, infected cell *p*, or the external force of infection *ext*—are determined by the transmission rate (β) between the infected and susceptible unit type, the relative infectivity of the infected unit (ψ) given its forest coverage (for cells) or temporal period (for domestic pig units), and indicator functions (1)) for both whether herd *i* or cell *p* is infectious and whether the infectious herd or cell is able to infect the susceptible unit (determined by a distance threshold for transmission between herds, adjacency for transmission between cells, and the herd being contained within the cell for transmission from herds to cells or cells to herds). Transmission between herds was further informed by a step function bound at 20 km for the distance between infectious herd *i* and susceptible herd *j* (*d*_*i,j*_), though parametric distance kernels (δ) were also tested. When assessing frequency-dependent transmission between herds and from cells to herds, the respective forces of infection were additionally divided across the number of herds over whom the infectious pressure was distributed.

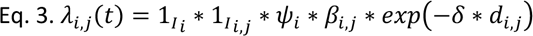

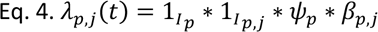

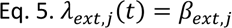

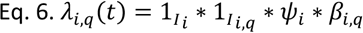

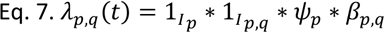

Transitions between states were stochastically modelled and based on a tau-leap algorithm (10). The probability of passive detection—indicated by the transition from the *Iu* to *Id* state—was considered as an exponentially-distributed function of the detection rate *σ,* with a probability of detection at a given time step defined as *p*_*det.passive*_ = 1 − *exp*(−σ). To account for active surveillance upon domestic pig units, the detection probability for domestic pig units was further modified by parameter *ζ*, a multiplicative term strictly larger than 1 to account for the increase of the detection rate within a surveillance zone (defined as a 10 km radius around a detected infectious herd, see supplemental material: determination of model parameters) at that time step and defined as: *p*_*det.passive*_ = 1 − exp (−(σ ∗ ζ)) (79). Assuming that all infections among domestic pig units were eventually detected, the probability of recovery from infection was governed by the exponentially-distributed recovery rate gamma, defined as: *p*_*rec*_ = 1 − *exp*(−γ). Given current understandings of carcass persistence, imperfect carcass detection, and environmental contamination, and the duration of the study period, cells were assumed to not become recovered following infection (44, 80).

Simulations were initialized via the infection of the two cells in Tulcea County which contained the two first detected wild boar cases in Romania, 8 weeks prior to their detection, as reported on 12 and 16 June 2018.

### Model selection, parameterization and calibration

Twenty-four models were constructed to reflect unknown dynamics that could not be elucidated from the observed outbreak data. Herd-to-herd transmission was modelled as either frequency or density dependent, with density dependence modelled through either a step function where infectious pressure is equally exerted on all units within the infectious area, or a transmission kernel where infectious pressure scales with distance from the infectious unit, both bounded at 20 km. Cell-to-herd transmission was also modelled as either frequency or density dependent transmission, as 30% of the cells with villages contained more than one village. For cell-to-cell transmission, as all cells but those on the edge have the same number of neighbors, frequency or density dependence would be an almost-null distinction. Instead, neighbor- order adjacency was assessed, considering either first order adjacency for infectious pressure on an infectious cell’s immediate neighbors, or second order adjacency, to also include once-removed neighbors. Lastly, cells highly-suitable to wild boar were defined as either those cells with adequate forest coverage (15% or greater) or cells with adequate forest coverage plus the immediately adjacent cells. This alternative formulation was motivated by the fact that, in the observed data, though the majority of wild boar cases were discovered in forested cells (62%, see supplemental material: epidemic analysis), of the cases outside of forests, 75% were in cells immediately adjacent to forested cells.

Of the 15 parameters in the model, four were able to be informed from available data in the literature or observed data (Table 1). The detection rate for herds was estimated at 1/3 weeks^-1^ based on analysis of ASF detection in the Russian Federation (81), and set at 1/8 weeks^-1^ for cells based on estimates for a large wild boar population (82). The recovery rate for domestic pig units was defined per county per eight-week period and calculated from the observed data.

The remaining 10 parameters were estimated through adaptive population Monte Carlo (APMC) approximate Bayesian computation (ABC) (83). Sequential ABC methods guide the selection of parameter values (particles) through the comparison of summary statistics that describe the observed and simulated data. Summary statistics reflected the spatial and temporal distribution of detected outbreaks, using quarterly incidence per county and per entire study region for domestic pig units and wild boar cells, for a total of 56 summary statistics. APMC includes a weighted particle filtering methodology combined with an algorithm for auto-determining the rejection tolerance level after each generation of particle selection, to aid in convergence to the final parameter distributions. Relying on non-informative priors, particles to comprise the initial parameter sets of the estimated model parameters were drawn from uniform distributions. Following a round of 500 model simulations, weights were applied to each particle based on the resulting distance between simulated and observed summary statistics. The best-fitting fifty percent of particles were retained, and another 250 particles were computed. This process repeated itself until the proportion of newly generated particles that satisfied the preset final tolerance limit (set at 0.05) was reached. The tolerance level specifies the acceptable distance of a particle from the data, and through this algorithm a decreasing sequence of tolerance levels were used which directed particle sampling towards a high-likelihood parameter space, thus avoiding unnecessary time spent throughout the entire parameter field. At this limit, additional simulations would have limited effect on the final posterior distributions.

Model performances in reproducing the overall infection dynamics were compared through the mean squared error (MSE) of median weekly incidence for both herds and cells at the level of the entire study region.

### Alternative management scenarios

Alternative control strategies were chosen to consider improvements in either case detection or case prevention, being evaluated through the end effects on total epidemic size and relative host contribution. Improving passive surveillance among domestic pig units was simulated through reducing the average duration of the undetected-infectious period from three to two weeks. Reactive culling of an entire village following case detection in that village was simulated through forcing an average infectious period duration of one week across all villages. Preventive culling of domestic pig units following detection of a nearby wild boar case was simulated by transiting domestic pig units in an infectious-detected cell to the R state upon cell case detection. Lastly, environmental sanitation through wild boar carcass clearance was simulated by having cells progress to the recovered state following a four-week infectious-detected period instead of remaining in the infectious state. Alternate scenario outcomes were compared to baseline results through Bayesian estimation, comparing 95% highest density intervals (HDI) of Bayesian posterior probabilities: that is, the highest density interval of the distribution of the posterior values that produced the observed outcome, analogous to confidence intervals from null-hypothesis significance testing.

Data compilation, model implementation, parameterization and analysis were performed in R statistical software, version 4.1.3 “One Push-Up” (84). Model fitting was done through the package easyABC, and Bayesian estimation for comparison of alternative control strategies was conducted with the BEST package (85, 86). Scripts and data are publicly available via Zenodo (87).

## Acknowledgments

The authors would like to thank Inaporc and Agreenium for funding this research, the Romanian veterinary authorities for their field expertise, and the INRAE MIGALE bioinformatics facility (88) for providing computing and storage resources.

