## Supplemental material for "A multi-host mechanistic model of African swine fever emergence and control in Romania"

### Supporting Information for: A multi-host mechanistic model of African swine fever emergence and control in Romania

*Brandon H. Hayes

Additional information is provided on epidemic analysis, the determination of model parameters, infection state selection, and uncertainty analysis.

**Epidemic analysis**

The epizootic was analysed spatially and temporally by host (domestic pig or wild boar) and domestic pig herd type (village or industrial site). Raw case data was spatially and temporally aggregated to reflect the epidemiological units (village, industrial farm, or habitat cell) and time scale (weekly) of the model. Temporally, cases were aggregated to the ISO-standard week. Based on expert opinion solicited from local veterinary officials, villages were defined as continuously infectious as long as successive outbreaks were declared less than two weeks apart. While the observed control strategies result in an internal diversity within the epidemiological unit of the village, this facet was addressed through considering the infectious status of a village en-masse from first to last case detection per outbreak. Epidemiological units were examined through frequency measurements (prevalence, incidence, and cumulative incidence) to inform model development.

A total of 1128 cases were reported over the study period. The vast majority of cases were seen among backyard farms in villages (n=1001), along with 17 cases among industrial sites and 110 among wild boar. These cases were aggregated by epidemiological unit (village, industrial site, or cell) into 393 outbreaks, with an outbreak defined as a group of cases with less than two weeks between each case, per expert opinion solicited from local veterinary officials (1). Found carcasses comprised 63% (n=69) of wild boar cases, while 29% (n=32) were among wild boar killed during hunting (8% of cases did not provide a means of capture). Though only 5% (n=65) of cells contained wild boar cases, 62% (n=68) of wild boar cases were discovered in cells with forest coverage above the 15% habitat threshold.

Of the 313 villages that experienced outbreaks, 13% (n=40) saw a resurgence of ASF over the model period. The median time between reinfection events was 5 weeks (range: 3–11), with 85% of villages having only 1 reinfection occurrence. Most reinfection events occurred between weeks 16 and 24, Reinfection events were not evenly distributed amongst all counties, as most of the villages that experienced reinfection were located in Tulcea (n=17) or Braila (n=14) counties. Further, of the villages that experienced two or three reinfections, all were located in either Tulcea (n=4) or Braila (n=2) county.

**Determination of model parameters**

Individual village infectious periods were calculated as the time in weeks between the first and last case in an outbreak, with an outbreak defined as subsequent cases with less than 2 weeks between detection. Spatiotemporal heterogeneity was identified among village infectious periods, with quarterly differences both within and between counties (Table S1).

Surveillance zones were included to reflect observed control measures. Per Romanian regulation and in accordance with EU directives, following the detection of an infected pig farm, the affected herd was culled, movement restrictions of people, animals and products were enacted, and a 10 km surveillance zone was established around the culled herd which would be kept in place for 4 weeks (1, 2). If no additional infectious holdings were detected during this period, the surveillance zone would be removed. While an industrial site would be culled in entirety, backyard farms within a village were considered individually and neighboring backyard herds to the infected herd were not automatically culled, allowing several backyard farms to be reported successively over periods of several days or weeks.

**Infection state selection**

The infection models consisted of combinations of four states for each epidemiological unit to transit through—susceptible (S), infectious undetected ($I_u$), infectious detected ($I_d$), and recovered R). Two models were tested for each component, leading to four model combinations. Villages and industrial sites were considered to either become perpetually recovered or, following a two-week refractory period, re-susceptible to infection. Wild boar cells were considered to either be continuously infectious until the end of the study period or able to become re-susceptible following their infectious period. When combined, this yielded models of $SI_uI_dR$-$SI_uI_d$, $SI_uI_dRS$-$SI_uI_d$, $SI_uI_dR$-$SI_uI_dS$, and $SI_uI_dRS$-$SI_uI_dS$ state combinations. Infectious periods were calculated per county from observed data for each of the epidemiological unit classes. Model selection among infection state combinations was performed through calculating the mean squared error (MSE) of median weekly incidence and cumulative incidence for domestic pig units and for wild boar cells, respectively, for each model component.

To further inform transmission dynamics among wild boar cells, having assumed home range size, cell-to-cell transmission of both first-order and second-order adjacency was modelled. Long range wild boar dispersal beyond the assumed 25 $km^2$ home range can occur secondary to hunting pressure and has been observed to be up to a mean of 45 km from the origin (3, 4). Following model selection among the four infection state combinations, one hundred simulations were performed with both first and second order adjacency transmission. Differences in modelled dynamics were determined through comparing posterior probabilities by calculating the 95% Highest Density Intervals (HDIs). This analysis was conducted through the BEST package in R (5).

Two sets of infection states—with or without re-susceptibility—were considered for each host unit in model development. With domestic pig units experiencing either $SI_uI_dR$ or $SI_uI_dRS$ states and cells experiencing either $SI_uI_d$ or $SI_uI_dS$ states, the assessed model infection state combinations yielded $SI_uI_dR$-$SI_uI_d$, $SI_uI_dR$-$SI_uI_dS$, $SI_uI_dRS$-$SI_uI_d$, and $SI_uI_dRS$-$SI_uI_dS$ models. Following parameterization of each model variant via ABC-SMC, the total MSE of the median weekly incidence for domestic pig units and median weekly cumulative incidence for wild boar cells was calculated. The models without re-susceptibility among wild boar cells ($SI_uI_dR$- $SI_uI_d$ and $SI_uI_dRS$-$SI_uI_d$) had the lowest MSEs of 271.1 and 239.1, respectively (Fig. S6). The goodness-of-fit for domestic pig units and wild boar cells were then examined independently. The model without re-susceptibility better approximated the observed data among domestic pigs, as a worse MSE was seen when reinfections were considered (47.3 for $SI_uI_dR$ compared to 70.84 for $SI_uI_dRS$). Further, reinfection events, having occurred in 40 (13%) of villages, were few and augmentation of the domestic pig model through considering reinfection was not supported by the data. The $SI_uI_dR$-$SI_uI_d$ model was selected as the final model for further analysis.

The non-reinfection model was also evaluated with cell-to-cell transmission of both first-order and second-order adjacency. Both models produced similar results, with differences in the mean weekly incidence of cells and villages of -0.164 (95% HDI: -0.394–0.0657) and 0.594 (95% HDI: -1.34–0.18), respectively. As there was no significant difference between the effect of first-order and second-order cell-to-cell transmission adjacency, to account for long-range dispersal, second-order adjacency was chosen in the model.

**Uncertainty Analysis**

An uncertainty analysis was performed on two key assumed parameters, to determine their effect on final epidemic size: the maximum local transmission distance between domestic pig units and the detection rate augmentation that occurs within surveillance zones through parameter zeta (ζ). Differences in outcomes from parameter perturbation were evaluated through comparing the 95% highest density intervals (95% HDI) between groups: both between the baseline parameter and its alternative, and between adjacent parameter values. This analysis was also performed through the BEST package in R (5).

The maximum distance under which local transmission could occur was assumed to be 20 km, which was informed through the median distance between villages. Preliminary analysis revealed median distances between villages ranging from 33.6 km (Interquartile range (IQR): 21.4–47.6) in Calarasi county to 46.3 km (IQR: 28.8–63.2) in Constanta. The median lower-bounds of the IQR for all villages in the landscape (22.8 km) was used to approximate the assumed maximum local transmission distance, which was rounded to 20 km. As our model used a power-law transmission function, the estimated kernel value itself (parameter δ,

Eq. 3 main text) was expected to play a far more influential role in guiding transmission than the maximum distance. However, the maximum distance was expected to play a larger role in dictating transmission when a step-function was considered, and alternative transmission limits of 10, 15, 25, and 30 km were explored.

Expectedly, epidemic size among both domestic pig units and wild boar cells was sensitive to this parameter (Fig. S7). Decreasing the maximum transmission distance from 20 km to 15 km and 10 km resulted in 52.5 (95% HDI: 33.7, 72.4) and 100.9 (95% HDI: 83.3,118.9) fewer cases among domestic pigs, and 19.6 (95% HDI: 10.8, 29.5) and 33.5 (95% HDI: 25.6, 43.1) fewer cases among wild boar (Table S2a). On average, each 5 km increase in maximum distance resulted in 43.6 additional domestic pig and 15 additional wild boar cases, though no significant increase in wild boar cases was seen when the maximum distance was increased from 25 km to 30 km.

The effect of the surveillance zone—that is, the relative increase in detection rate of ASF cases for units in the surveillance zone compared to those outside the zone, and given by the parameter zeta—was initially assumed to double the rate of detection compared to areas not under surveillance, and represents the effectiveness of the field activities on case detection. To evaluate the effect of this parameter, simulations were also conducted with surveillance zones having no additional effect, as well as increasing the relative detection rate by 1.5x, 2.5x, and 3x times the non-surveillance zone detection rate.

The increase in detection rate in surveillance zones resulted in significant decreases in epidemic size among both domestic pigs and wild boar for almost all scenarios (Fig. S8, Table S2b). Only when the surveillance zone effect was 1.5x the baseline was a non-significant effect on final epidemic size seen, and this was only in the wild compartment.

Greater surveillance zone effects resulted in greater decreases in final epidemic size, however when adjacent surveillance zone effect parameter values were compared, a significant decrease in final epidemic size among both hosts was only seen when the surveillance zone effect was increased from 1.5x to 2x (Table S2b). This suggests that smaller increases in the effect of the surveillance zones will likely have little or no effect on the final epidemic size, however larger increases (e.g. doubling or tripling the case detection rate in these zones) can have impactful effects on epidemic mitigation.

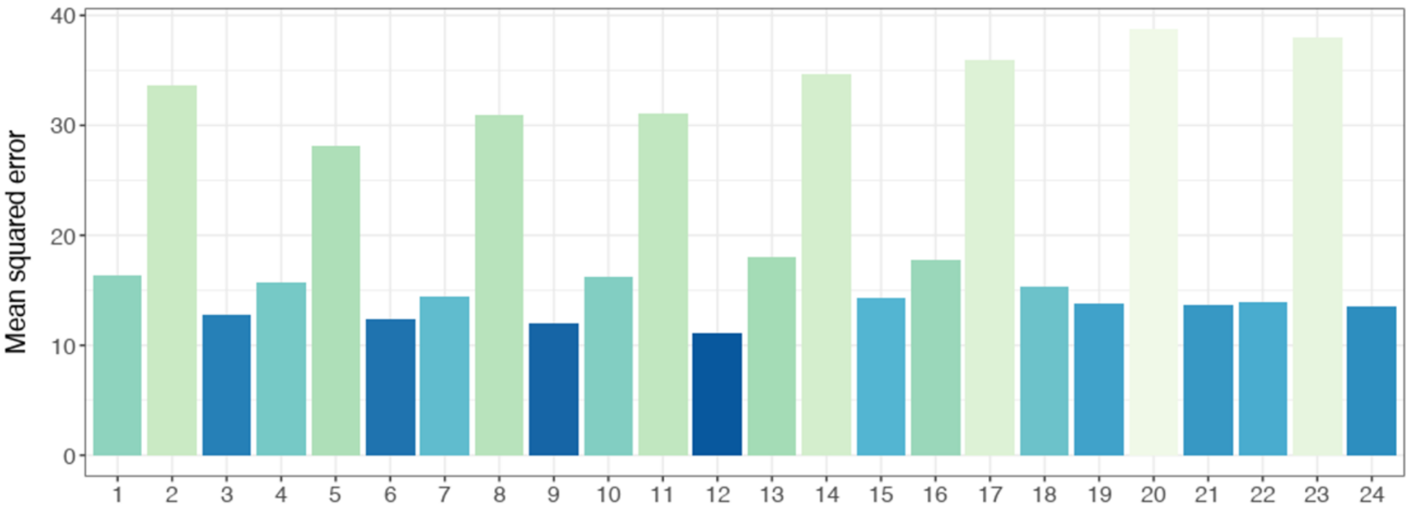


| | 1 | 2 | 3 | 4 | 5 | 6 | 7 | 8 | 9 | 10 | 11 | 12 | 13 | 14 | 15 | 16 | 17 | 18 | 19 | 20 | 21 | 22 | 23 | 24 |
|---|---|---|---|---|---|---|---|---|---|---|---|---|---|---|---|---|---|---|---|---|---|---|---|---|
| dp-dp | DK | DS | FS | DK | DS | FS | DK | DS | FS | DK | DS | FS | DK | DS | FS | DK | DS | FS | DK | DS | FS | DK | DS | FS |
| wb-dp | D | D | D | D | F | F | F | F | D | D | D | D | F | F | F | F | D | D | D | D | F | F | F | F |
| wb-wb | 1 | 1 | 1 | 1 | 1 | 1 | 2 | 2 | 2 | 2 | 2 | 2 | 1 | 1 | 1 | 1 | 1 | 1 | 2 | 2 | 2 | 2 | 2 | 2 |
| for-neigh. | F | F | F | F | F | F | F | F | F | F | F | F | T | T | T | T | T | T | T | T | T | T | T | T |

**Fig. S1.** Comparison of fit by mean squared error (MSE) of the 24 evaluated model combinations. Twenty-four models were constructed on combinations of between-host and within-host transmission assumptions, and compared through MSE of median weekly incidence among herds and cells. Below each model number (1–24) are indicators for the characteristics of the given model. For transmission between domestic pig units (dp-dp), the transmission mechanism was either density-dependent with a distance kernel (DK), density-dependent with a step function (DS) or frequency-dependent with a step function (FS). For transmission from wild boar cells to domestic pig herds (wb-dp), the evaluated transmission functions were of density-dependence (D) and frequency-dependence (F). For transmission between wild boar cells (wb-wb), either first (1) or second (2) order adjacency between cells was considered for potential transmission distances. Lastly, both cells with sufficient forest coverage and cells immediately adjacent to cells with sufficient forest coverage were considered possibilities for suitable wild boar habitat. To determine which assumption better-represented the data, cells adjacent to forested cells (for-neigh.) were either excluded (F) or included (T) as sufficient wild boar habitat.

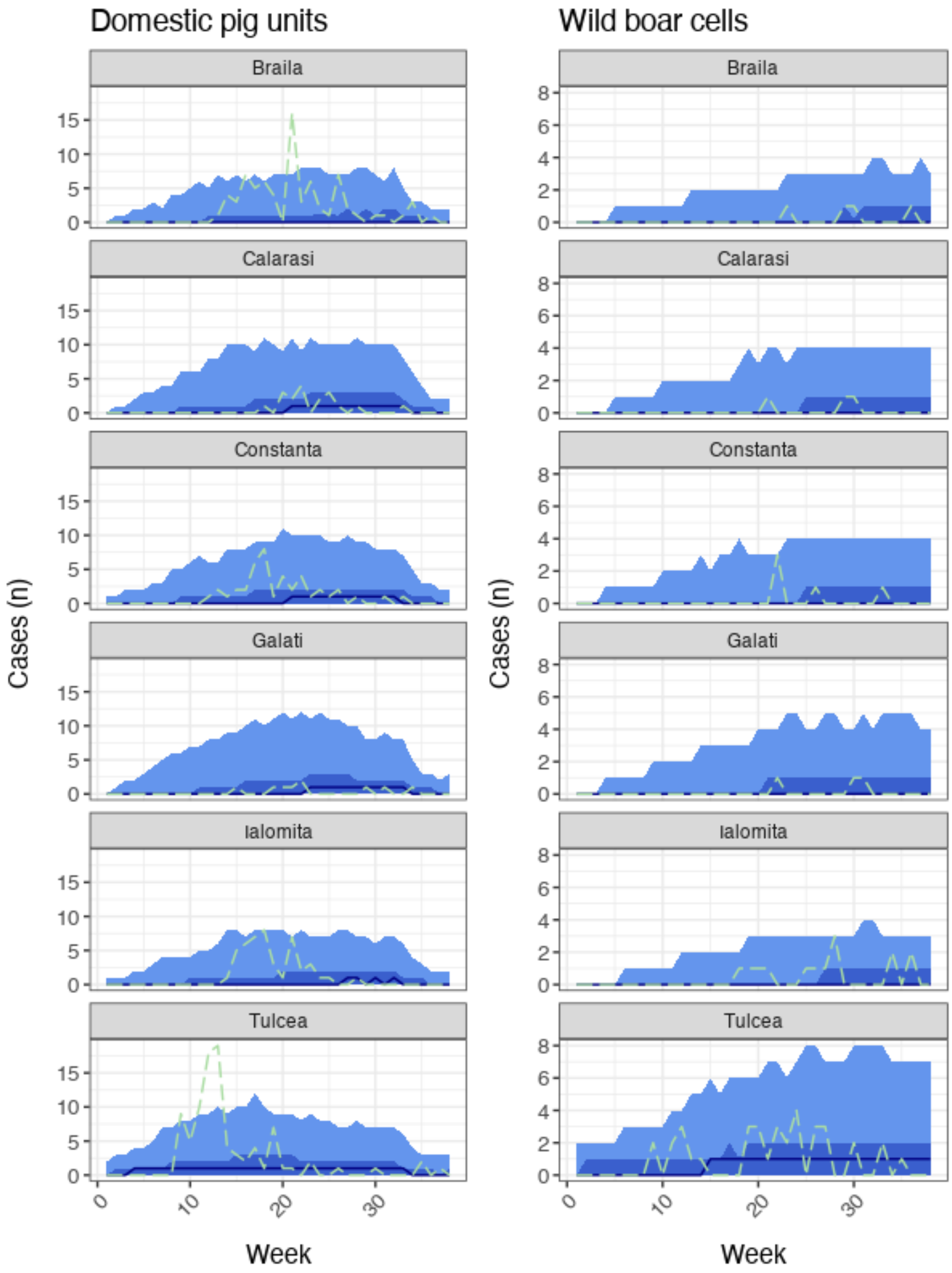


**Fig. S2.** Spatially-explicit (by county) weekly incidence trajectories of simulated epidemics. The light blue ribbon indicates the 99% credible interval—that is, a 99% probability the true epidemic incidence estimate lies within this region—while the darker blue ribbon indicates the 50% credible interval, the darkest blue line the simulated median, and the dashed green line the trajectory of the observed epidemic.

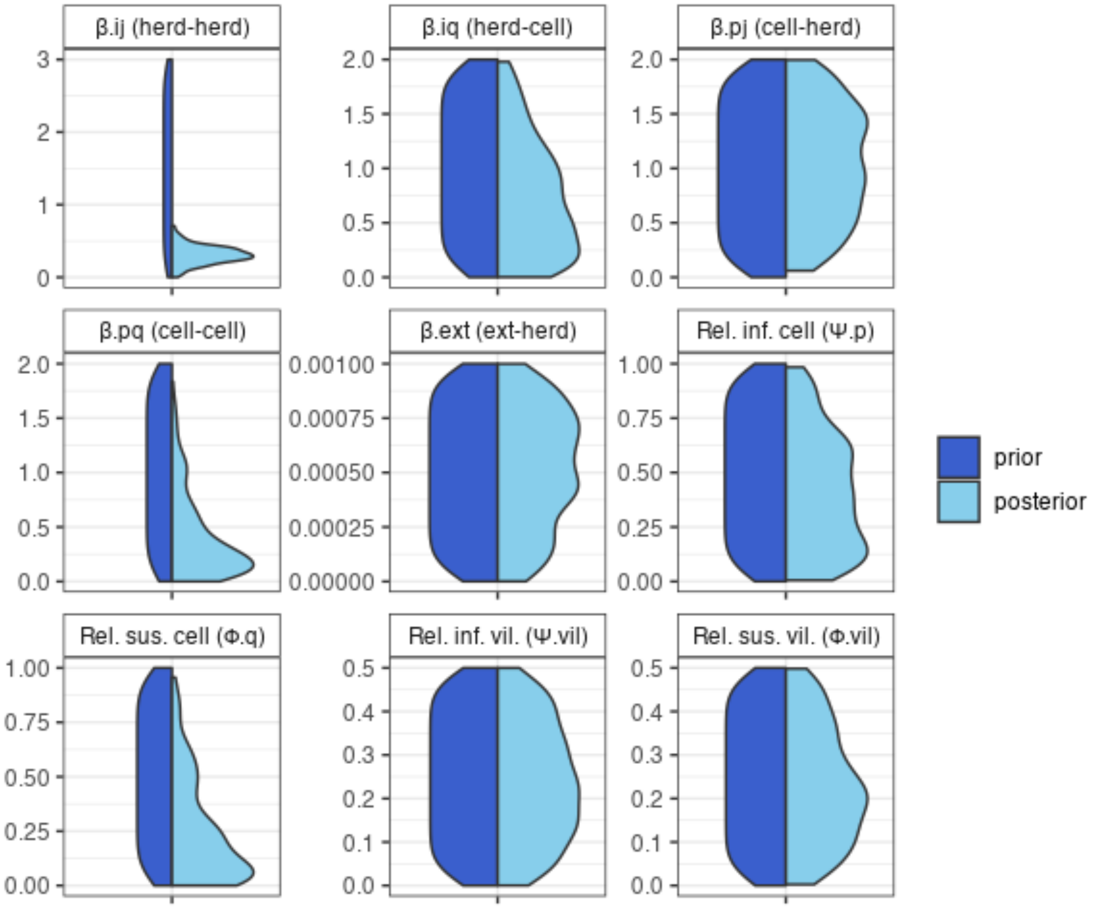


**Fig. S3.** Prior and posterior distributions of estimated model parameters. Uniform priors (dark blue) were used for all estimated model parameters. Y-axis units are dependent on the estimated parameter, with transmission rates (betas) in infections per week, and relative infectivity and susceptibility (psi and phi) in percent.

#### Domestic pig to wild boar transmission removed

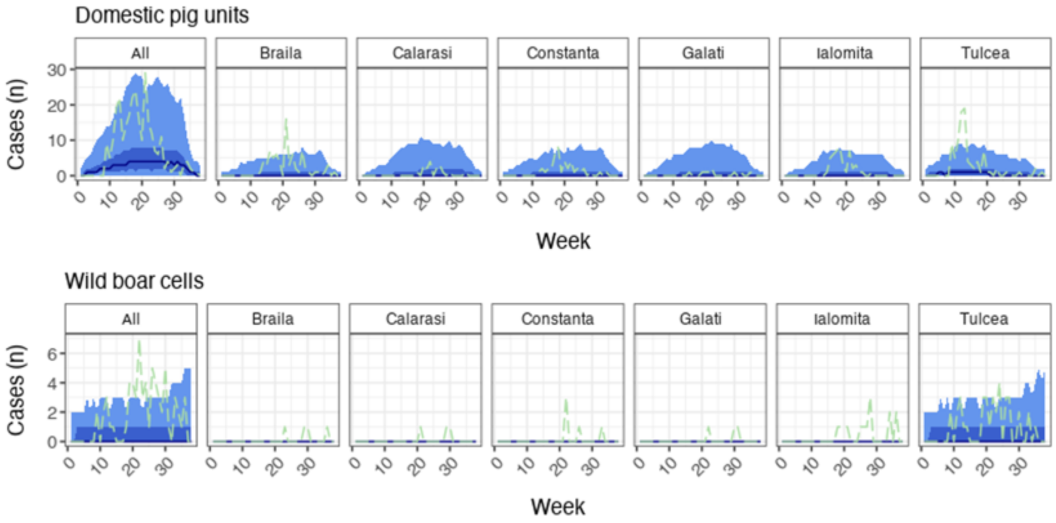


#### Wild boar to domestic pig transmission removed

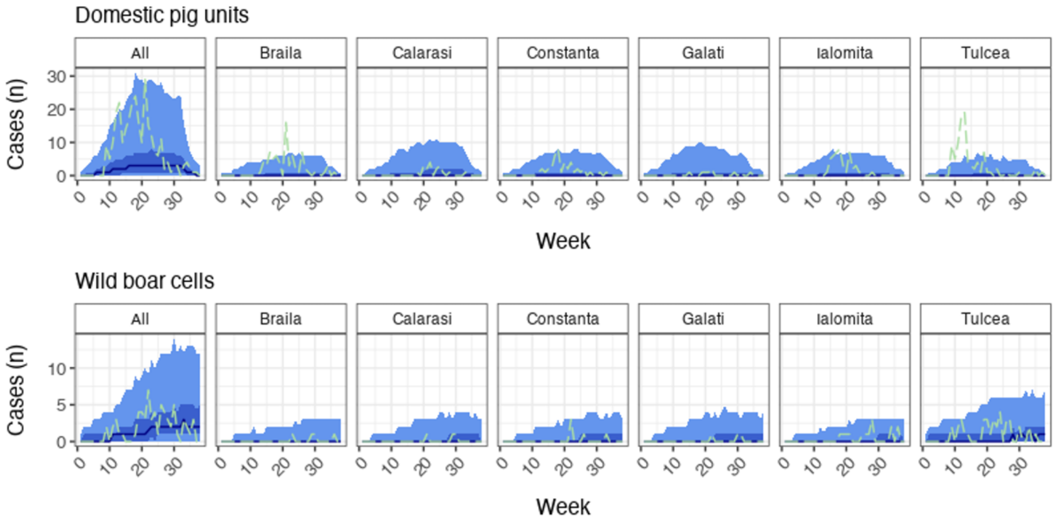


**Fig. S4.** Spatiotemporal dynamics with removal of single interhost transmission pathway. When interhost transmission from domestic pig units to wild boar cells was removed (top plot pair) from the model dynamic, the observed wild boar incidence was unable to be replicated, either per-county or en-masse. Conversely, when interhost transmission from wild boar cells to domestic pig units was removed (bottom plot pair), the epidemic trajectories among domestic pig units were reduced though the epidemic continued to be observed in both compartments. The light blue ribbon indicates the 99% credible interval, the darker blue ribbon the 50% credible interval, the darkest blue line the simulated median, and the dashed green line the observed data.

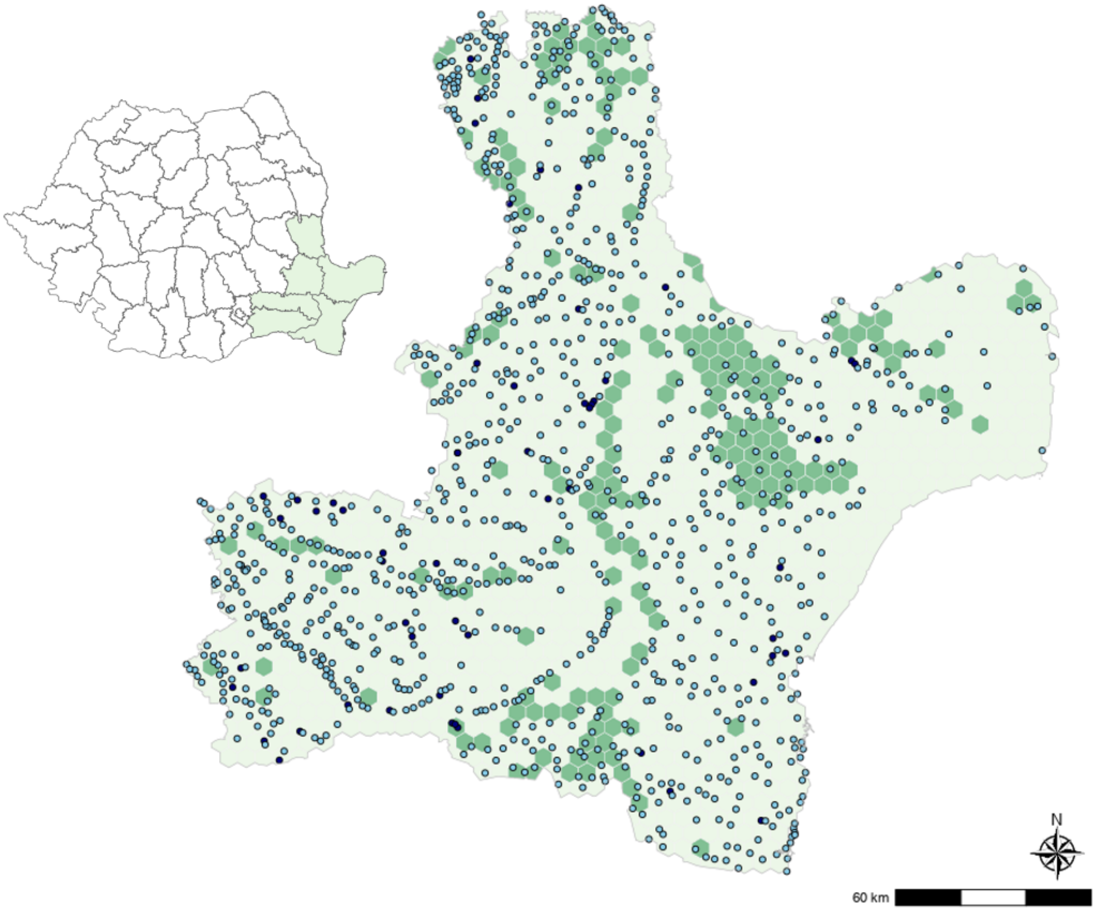


**Fig. S5.** Spatial visualization of model landscape. The region of interest was the area of initial epidemic spread, the six southeastern-most counties of Romania. Among domestic pig units, villages are depicted as light blue circles and industrial sites as dark blue circles. For determining suitable wild boar habitat, 25 km$^2$ hexagons were partitioned by percent forest coverage. Light green hexagons represent areas with less than 15% forest coverage and dark green hexagons represent suitable habitat areas containing 15% or more forest coverage.

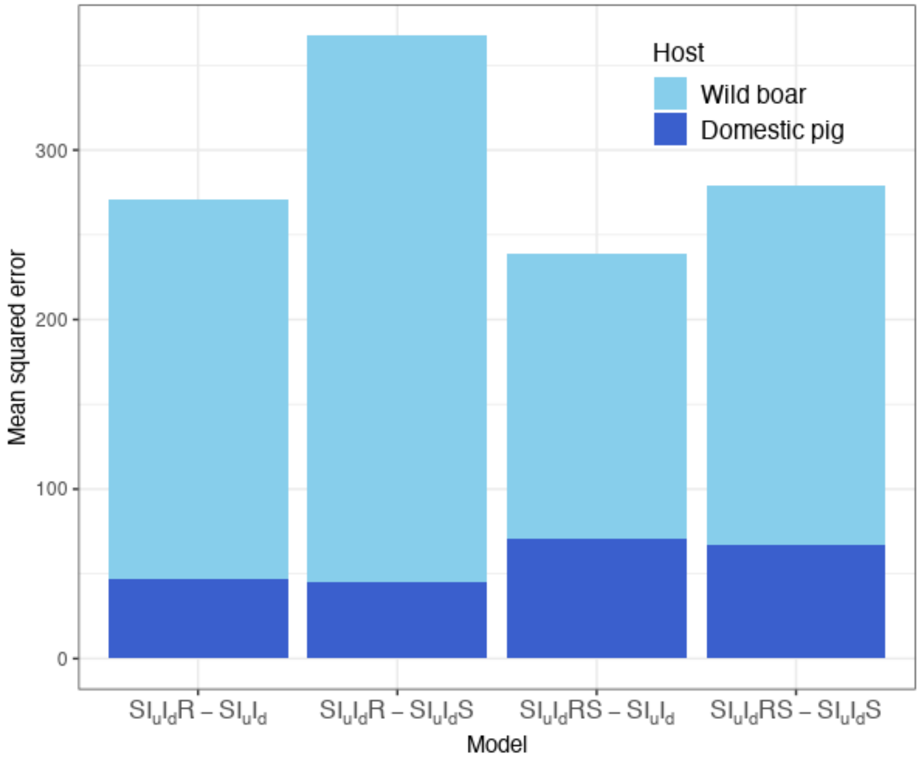


**Fig. S6.** Mean squared error (MSE) for infection-state model variants. Median weekly incidence for domestic pig units and cumulative incidence for wild boar cells was used to calculate the MSE for all infection-state model variants. Two models were tested for each component, yielding four model combinations. Domestic pig units (villages and industrial sites) were modelled through either $SI_uI_dR$ or $SI_uI_dRS$ states, and wild boar cells were modeled with $SI_uI_d$ or $SI_uI_dS$ infection states.

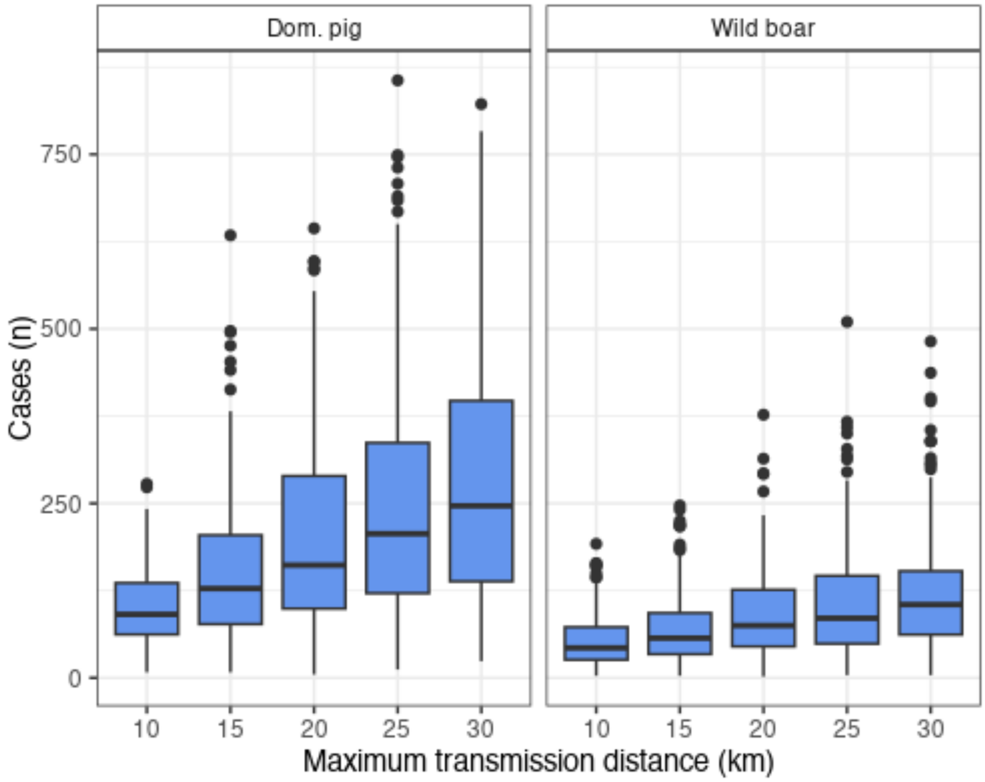


**Fig. S7.** Effect of maximum transmission distance on final epidemic size, 250 simulations per scenario. Each 5 km increase in maximum distance resulted in an average of 43.6 additional domestic pig cases and 15 additional wild boar cases.

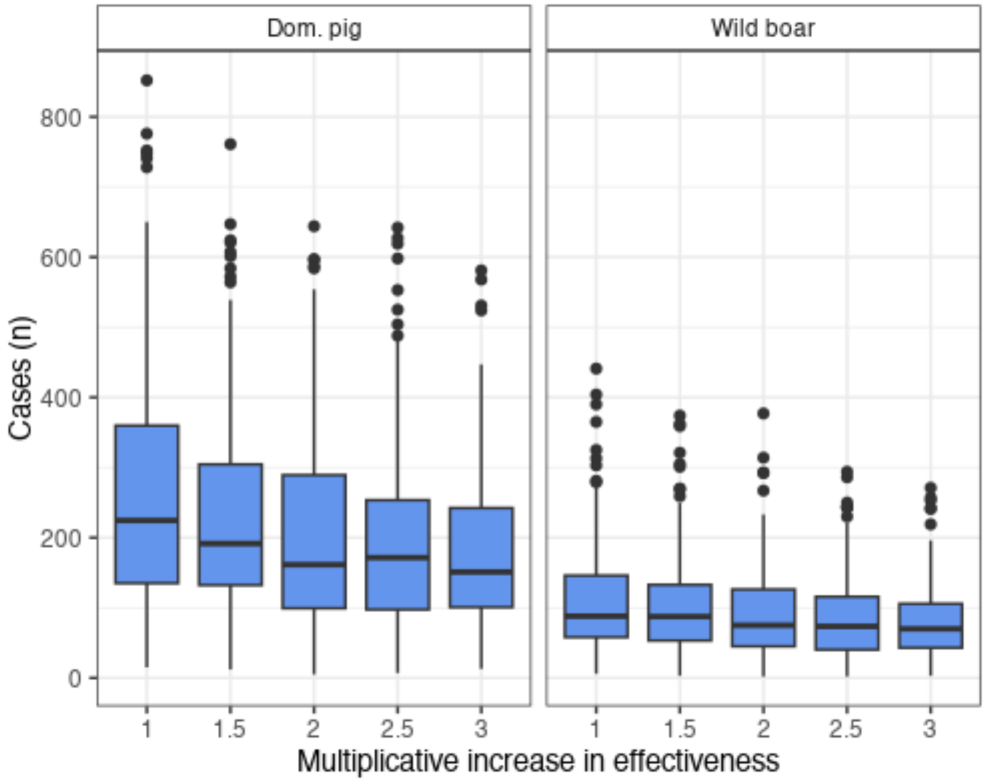


**Fig. S8.** Effect of surveillance zone strength on final epidemic size, 250 simulations per scenario. Compared to the estimated epidemic size had no surveillance zones been enacted, significant reductions were seen for all parameter values except among wild boar for when the surveillance zone was only 1.5x as effective. When adjacent surveillance zone effect values were compared, significant differences were seen only among domestic pigs when increasing the effectiveness from 1 to 1.5x and from 1.5x to 2x, and only among wild boar when increasing the effectiveness from 1.5x to 2x.

**Table S1.** Spatiotemporally explicit recovery rate values ($\gamma_{i.vil}$)

| County | Quarter | Value (weeks$^{-1}$) | County | Quarter | Value (weeks$^{-1}$) |
|---|---|---|---|---|---|
| Braila | 1 | 0.787 | Galati | 1 | 0.500 |
| | 2 | 0.840 | | 2 | 0.5 |
| | 3 | 0.893 | | 3 | 1 |
| | 4 | 1 | | 4 | 1 |
| Calarasi | 1 | 0.344 | Ialomita | 1 | 0.500 |
| | 2 | 0.344 | | 2 | 0.741 |
| | 3 | 0.350 | | 3 | 1 |
| | 4 | 1 | | 4 | 1 |
| Constanta | 1 | 0.609 | Tulcea | 1 | 0.413 |
| | 2 | 0.769 | | 2 | 0.763 |
| | 3 | 1 | | 3 | 1 |
| | 4 | 1 | | 4 | 1 |

**Table S2a. Effects of maximum local transmission distance on final epidemic size**

| Maximum distance value (km) | Median change in epidemic size (95% HDI) | |
|---|---|---|
| | Domestic pig units | Wild boar cells |
| From baseline (20) to: | | |
| 10 | -100.9 (-118.9, -83.3) | -33.5 (-43.1, -25.6) |
| 15 | -52.5 (-72.4, -33.7) | -19.6 (-29.5, -10.8) |
| 25 | 39.7 (12.7, 64.3) | 12.9 (2.9, 25.2) |
| 30 | 78.3 (50, 105.7) | 26.5 (13.7, 38.4) |
| From adjacent value: | | |
| 10 → 15 | 43 .9 (29.5, 56.3) | 14.3 (6.8, 20.6) |
| 15 → 20 | 51.1 (31.9, 72.7) | 19.7 (10.9, 28) |
| 20 → 25 | 40.6 (14.3, 67.8) | 12.8 (2, 23.6) |
| 25 → 30 | 37.7 (8.9, 65.2) | 13.1 (-1.2, 24.4)* |

**Table S2b. Effects of surveillance zone multiplier on final epidemic size**

| Surv. zone multiplier increase | Median change in epidemic size (95% HDI) | |
|---|---|---|
| | Domestic pig units | Wild boar cells |
| From baseline (1.0) to: | | |
| 1.5 | -33.7 (-59.2, -9.1) | -5.3 (-16.2, 6.8)* |
| 2.0 | -60.5 (-92.5, -36.4) | -16.4 (-27.4, -5.5) |
| 2.5 | -71 (-96.6, -44.7) | -19.2 (-30.1, -9.1) |
| 3.0 | -84.5 (-106.6, -57.6) | -25.1 (-34.2, -13.7) |
| From adjacent multiplier: | | |
| 1.5 → 2.0 | -26.1 (-51.6, -3.7) | -10.8 (-21.4, -0.5) |
| 2.0 → 2.5 | -12 (-33.1, 10.9)* | -3.3 (-14, 6.3)* |
| 2.5 → 3.0 | -12.7 (-32, 7.5)* | -5 (-13.3, 4.2)* |

*95% HDI contains zero, meaning the parameter perturbation likely had no effect on the observed difference in outcomes of transmission between the compared scenarios.
